# DsrMKJOP is the terminal reductase complex in anaerobic sulfate respiration

**DOI:** 10.1101/2023.08.03.551783

**Authors:** Ana C. C. Barbosa, Sofia S. Venceslau, Inês A. C. Pereira

## Abstract

Microbial dissimilatory sulfate reduction is a key process in the Earth biogeochemical sulfur cycle. In spite of its importance to the sulfur and carbon cycles, industrial processes and human health, it is still not clear how reduction of sulfate to sulfide is coupled to energy conservation. A central step in the pathway is the reduction of sulfite by the DsrAB dissimilatory sulfite reductase, which leads to the production of a DsrC-trisulfide. A membrane-bound complex, DsrMKJOP, is present in most organisms that have DsrAB and DsrC, and its involvement in energy conservation has been inferred from sequence analysis, but its precise function was so far not determined. Here, we present studies revealing that the DsrMKJOP complex of the sulfate reducer *Archaeoglobus fulgidus* works as a menadiol:DsrC-trisulfide oxidoreductase. Our results reveal a close interaction between the DsrC-trisulfide and the DsrMKJOP complex and show that electrons from the quinone pool reduce consecutively the DsrM hemes *b*, the DsrK noncubane [4Fe-4S]^3+/2+^ catalytic center, and finally the DsrC-trisulfide with concomitant release of sulfide. These results clarify the role of this widespread respiratory membrane complex and indicate that DsrMKJOP will provide the missing link to energy conservation by generating a *proton motive force* across the membrane in the last step of dissimilatory sulfate reduction.

**Significance Statement:** Dissimilatory sulfate reduction (DSR) is a vital microbial process in anoxic environments, namely in sulfate-rich marine sediments that harbor a vast microbial ecosystem. DSR drives the global biogeochemical sulfur cycle and is crucial in remineralization of organic matter on the seafloor. It also has huge environmental impact by preventing release of the greenhouse gas methane from these sediments, through its oxidation coupled to sulfate reduction. Despite its high ecological importance, it is still not clear how microorganisms derive energy to grow through DSR. Here, we disclose the physiological function of a widespread membrane complex in DSR, showing it acts as the terminal reductase in the respiratory chain and providing important insights into how sulfate/sulfite reduction is linked to energy conservation.

## INTRODUCTION

Sulfur is a key element for life, mainly due to its chemical versatility and ability to switch between several biologically relevant oxidation states, ranging from -2, as in hydrogen sulfide (H_2_S), up to +6, as in sulfate (SO_4_^2-^). A critically important microbial process in the biogeochemical sulfur cycle is dissimilatory sulfate reduction (DSR) driven by anaerobic bacteria and archaea, in which the reduction of sulfate to hydrogen sulfide is linked to energy conservation (1, 2). This process has particular impact in marine sediments, accounting for up to 50% of carbon mineralization in the sea floor (3), and preventing methane emissions through its contribution to anaerobic methane oxidation (4, 5). Despite being a very ancient bioenergetic pathway (6), it is still not clear how sulfate reducers are able to conserve energy during sulfate respiration. In DSR, a central step is the reduction of sulfite, performed by the dissimilatory sulfite reductase DsrAB and its physiological partner DsrC, which mediate a four-electron reduction of sulfite to zero-valent sulfur in the form of a trisulfide formed between two conserved cysteines of DsrC (7–9). The reduction of the DsrC-trisulfide to hydrogen sulfide is thought to derive electrons from the quinone pool via the membrane-bound DsrMKJOP complex, pointing to a critical role of this complex in energy conservation (7, 10, 11). The Dsr proteins, namely DsrAB, DsrC and DsrMKJOP, are widespread in the microbial world, being also present in phototrophic and chemotrophic sulfur-oxidizing organisms where they are involved in sulfite oxidation (12, 13), and in many organisms capable of sulfur, thiosulfate, sulfite or organosulfonates reduction, and in sulfur disproportionators (14, 15). However, there is still no concrete evidence demonstrating the physiological function of the DsrMKJOP complex.

The DsrMKJOP complex was first isolated and characterized from *Archaeoglobus fulgidus* (16), then from *Desulfovibrio desulfuricans* (11), and later from *Allochromatium vinosum* (17), where this complex has been shown to be essential for sulfur globule oxidation (18). This transmembrane complex contains two periplasmic subunits (DsrJ and DsrO), two integral membrane subunits (DsrM and DsrP) and a cytoplasmic subunit (DsrK), and appears to be composed of a combination of two modules, DsrMK and DsrJOP (19). The MK module is homologous to the membrane-bound HdrED heterodisulfide reductase (Hdr) from methanogens (20–22), comprising a quinone-interacting membrane diheme cytochrome *b* DsrM, with both hemes displaying His/His coordination, and an iron-sulfur subunit DsrK predicted to have two canonical [4Fe-4S]^2+/1+^ clusters, and one noncubane [4Fe-4S]^3+/2+^ cluster (11). The noncubane cluster is predicted on the basis of a conserved five-cysteine motif CX_n_CCGX_n_CX_2_C (the CCG motif) (23), which was identified to bind the catalytic noncubane cluster for heterodisulfide reduction in the homologous HdrD (16). The DsrJOP module contains DsrP, an integral membrane subunit with ten predicted transmembrane helices, and DsrO, an iron-sulfur subunit predicted to coordinate three to four [4Fe-4S]^2+/1+^ clusters, which together resemble the quinone-interacting NrfCD module widespread in bacterial oxidoreductases (24). DsrJ is a membrane-anchored triheme cytochrome *c*, where each heme is proposed to have distinct axial coordination: His/His, His/Met and His/Cys (11, 25). The His/Cys heme *c* coordination is very unusual but is also found in SoxXA (26), TdsA (27) and PufC (28), which are involved in thiosulfate oxidation and/or electron transfer, suggesting that DsrJ could have a catalytic role in sulfur redox chemistry. Notably, the DsrJOP module is absent in many Gram-positive (Firmicutes) and archaeal sulfate reducers (19), and so does not seem to be essential. In contrast, the DsrMK proteins are present in all organisms containing DsrAB and DsrC, often encoded in the same operon and sometimes with multiple copies in the genome, which suggests this is the minimum functional module of this respiratory complex (19). Based on its similarity to HdrED, the DsrMK module was proposed to catalyze the reduction of DsrC-trisulfide to hydrogen sulfide through the oxidation of the quinone pool (7, 10, 11, 29). However, the presence of the DsrJOP module suggests that there may be a more elaborate mechanism (7). Both DsrM and DsrP are homologous to quinone-interacting proteins, and the presence of two putative quinone-binding sites led to the suggestion that a quinone cycle between the DsrMK and DsrJOP modules may be present (10).

The X-ray structure of the soluble HdrABC-MvhAGD complex from the methanogenic archaeon *Methanothermococcus thermolithotrophicus*, comprising the MvhAGD hydrogenase and the HdrABC heterodisulfide reductase, provided the first structural insights into the two noncubane [4Fe-4S]^3+/2+^ clusters present in the active site of HdrB, and how they mediate the reduction of CoM-S-S-CoB heterodisulfide (30). Each noncubane cluster is ligated by five cysteines from the predicted CCG motif. There are two such motifs in both HdrB and HdrD, the catalytic subunits of the two different types of Hdrs in methanogens. Cleavage of the CoM-S-S-CoB disulfide bond has been suggested to involve the two noncubane clusters in a homolytic mechanism, along with binding of each substrate to one of the clusters (30). Remarkably, only one CCG motif is present in DsrK, raising the question of how this protein may operate with a single noncubane cluster. Recently, studies of the noncubane [4Fe-4S]^3+/2+^ cluster in the HdrB protein from *Desulfovibrio vulgaris* Hildenborough revealed that this cluster can adopt “closed” and “open” conformations corresponding to the substrate-free and substrate-bound forms, respectively (31), suggesting that cluster flexibility may be important for binding the DsrC-trisulfide in the active site of its homolog, DsrK.

Overall, unraveling the role of the DsrMKJOP complex and its link to DsrC-trisulfide reduction and corresponding energy conservation mechanism is a major challenge in understanding dissimilatory sulfur metabolism. In the present study, we provide evidence for the activity of the DsrMKJOP complex as a menadiol:DsrC-trisulfide oxidoreductase, clarifying the functional role of this membrane complex in the final step of the sulfate respiratory chain, where it is likely coupled to energy conservation.

## RESULTS AND DISCUSSION

### DsrMKJOP isolation and reduction with Menadiol

The DsrMKJOP transmembrane complex was purified with DDM as solubilizing detergent from membranes of the thermophilic archaeon *A. fulgidus,* by adapting the protocol previously reported by Mander *et al.* (16). The DsrMKJOP complex is the only heme-containing membrane complex in *A. fulgidus* (19), which facilitates its purification compared with other sulfate reducers, such as *D. desulfuricans* (11). In *A. fulgidus*, the predicted molecular mass for each subunit of the complex is 38.4 kDa (DsrM, locus tag: AF0501), 64.4 kDa (DsrK, AF0502), 16.7 kDa (DsrJ, AF0503), 30.5 kDa (DsrO, AF0499), and 44.2 kDa (DsrP, AF0500). The purified complex displayed four bands on a Tricine-SDS-PAGE gel (**Figure S1**) with apparent molecular masses of 61, 31, 28, and 17 kDa, in line with the report by Mander *et al.* (16). The 17 kDa band was identified as a cytochrome *c* (DsrJ) by heme-staining, and the other bands were assigned as DsrK (61 kDa), DsrM (31 kDa), DsrO (28 kDa), based on previous identifications (11, 16). As reported before, DsrP is not detected by SDS-PAGE (11, 16) because it is a highly hydrophobic subunit (ten predicted transmembrane helices), which hinders its migration in the gel.

The UV-visible spectrum of the as-purified DsrMKJOP (oxidized state) is dominated by absorption maxima at 280, 410 and 527 nm (**Figure 1A**), confirming the presence of hemes. Upon reduction, the Soret peak shifts to 420 nm, and the β- and α-bands are observed at 524 and 557 nm, respectively, which is in line with previously reported spectra of DsrMKJOP from *A. fulgidus* (16) and *D. desulfuricans* (11). The α-band is quite broad supporting the presence of hemes *b* and *c*. Addition of menadiol reduced 38% of the total hemes as compared with full reduction by dithionite. This corresponds to approximately two hemes reduced, and since hemes *b* typically have higher redox potentials than hemes *c,* this likely corresponds to the two predicted hemes *b* in the quinone-interacting DsrM membrane subunit. A similar result was reported by Pires *et al.* (11), where 40% of hemes belonging to the *D. desulfuricans* DsrMKJOP were also reduced upon incubation with menadiol.

**Figure 1.**
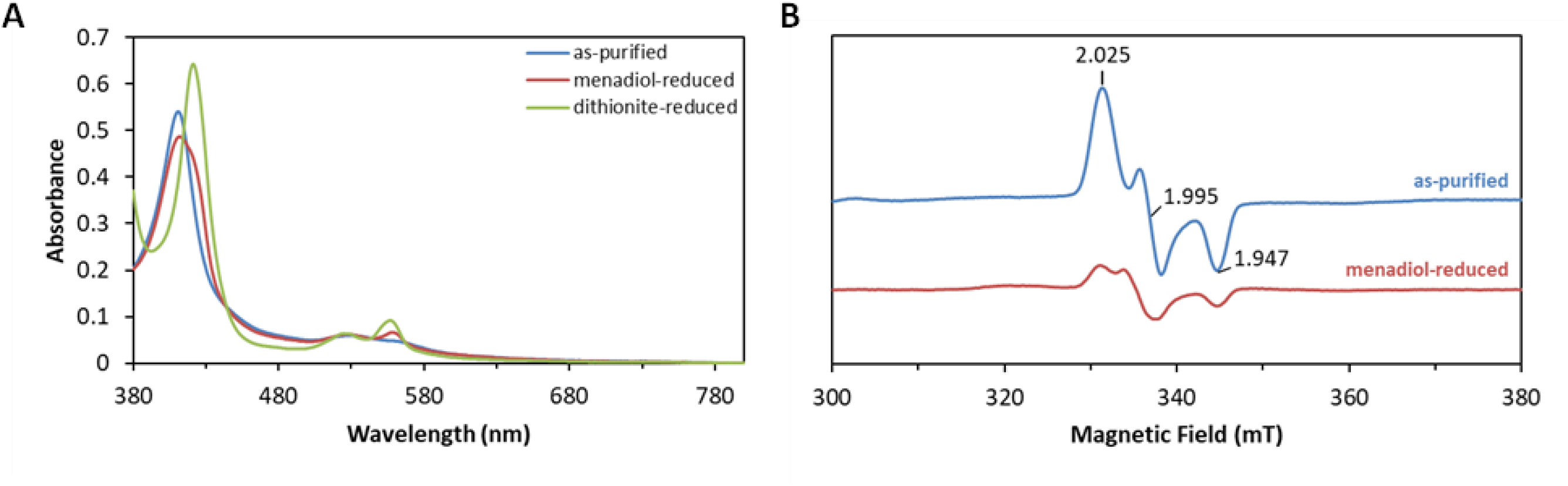
– UV-visible and EPR spectroscopy of the *A. fulgidus* DsrMKJOP complex. (A) UV-visible spectra of as-purified DsrMKJOP (blue), menadiol-reduced (red), and dithionite-reduced (green). (B) EPR spectra of the [4Fe-4S]^3+/2+^ cluster of the as-purified DsrMKJOP (blue), and upon menadiol reduction (red). *g*-values are marked in the spectrum of as-purified DsrMKJOP. All menadiol reduction spectra were obtained in a 1:500 protein to menadiol ratio.

The electron paramagnetic resonance (EPR) spectrum of the as-purified *A. fulgidus* DsrMKJOP shows an intense rhombic signal characteristic of a noncubane [4Fe-4S]^3+/2+^ cluster in the oxidized state, with *g*-values of 2.025, 1.995 and 1.947 (**Figure 1B**). This EPR signature was first observed in heterodisulfide reductases from methanogens (upon addition of HS-CoM or HS-CoB), with *g*-values of 2.013, 1.991 and 1.938 for HdrABC from *Methanothermobacter marburgensis* and 2.012, 1.993 and 1.946 for HdrED from *Methanosarcina barkeri* (41, 42), both of which possess two noncubane [4Fe-4S]^3+/2+^ catalytic centers involved in mediating heterodisulfide (CoM-S-S-CoB) reduction in two one-electron steps (22, 43). The previously reported *g*-values of *A. fulgidus* DsrMKJOP were 2.031, 1.994 and 1.941 (16), while in *D. desulfuricans* DsrMKJOP the *g*-values were 2.027, 1.991 and 1.943 (11). Given the similarity of DsrK to HdrD, the noncubane [4Fe-4S]^3+/2+^ cluster present in DsrK is likely involved in thiol/disulfide chemistry, suggesting that DsrK is the catalytic subunit of the DsrMKJOP complex. Upon incubation with menadiol under anaerobic conditions, the intensity of the rhombic signal decreased by 85%, revealing a pathway for electron transfer from the hemes *b* of DsrM to the noncubane [4Fe-4S]^3+/2+^ cluster of DsrK. The redox potential of the noncubane center in *A. fulgidus* DsrK was previously determined to be +90 mV (16), which agrees with its reduction by menadiol (midpoint potential of approx. -70 mV).

### The DsrC-trisulfide interacts strongly with DsrMKJOP

In *A. vinosum*, coelution upon protein purification suggested that DsrK interacts with DsrC (17). Following our identification of the DsrC-trisulfide as the product of sulfite reduction by DsrAB, we proposed that this is the actual substrate for the DsrMKJOP complex (7). To assess if there is a protein-protein interaction between the DsrC-trisulfide and the DsrMKJOP complex, surface plasmon resonance (SPR) was initially used. SPR is a highly sensitive technique for studying protein-protein interactions, without the need for labeling. The *A. fulgidus* DsrC-trisulfide was produced by *in vitro* enzymatic reaction using DsrAB from the same organism, as described in Santos *et al.* (7). The DsrC-trisulfide thus produced was immobilized on a nitrilotriacetic acid (NTA) sensor chip by the N-terminal His-tag, and the interaction was studied with increasing concentrations of DsrMKJOP complex (**Figure 2**). A strong interaction was observed between the two proteins, which could be detected even at low concentrations of the membrane complex. Steady-state conditions could not be achieved (even by increasing the association time), which means that no kinetic model could fit the experimental data, and so the association and dissociation rate constants could not be obtained. Nevertheless, an estimate for the equilibrium dissociation constant (K_D_) of 200 ± 27 nM was derived from duplicate sensorgrams using the maximum response reached. This K^D^ confirms a strong interaction between the DsrC-trisulfide and the DsrMKJOP complex (strong interactions are in the nanomolar range, or even lower as in the case of high-affinity antibodies), supporting the DsrC-trisulfide as a possible substrate for the membrane-bound DsrMKJOP complex. The SPR data were obtained at 25 °C, whereas a higher temperature (as used in the activity assays) should favor the association and dissociation rates of DsrC-trisulfide to DsrMKJOP complex since *A. fulgidus* is a thermophile, suggesting that the actual K_D_ at physiological temperatures may be somewhat different, and possibly even lower.

**Figure 2.**
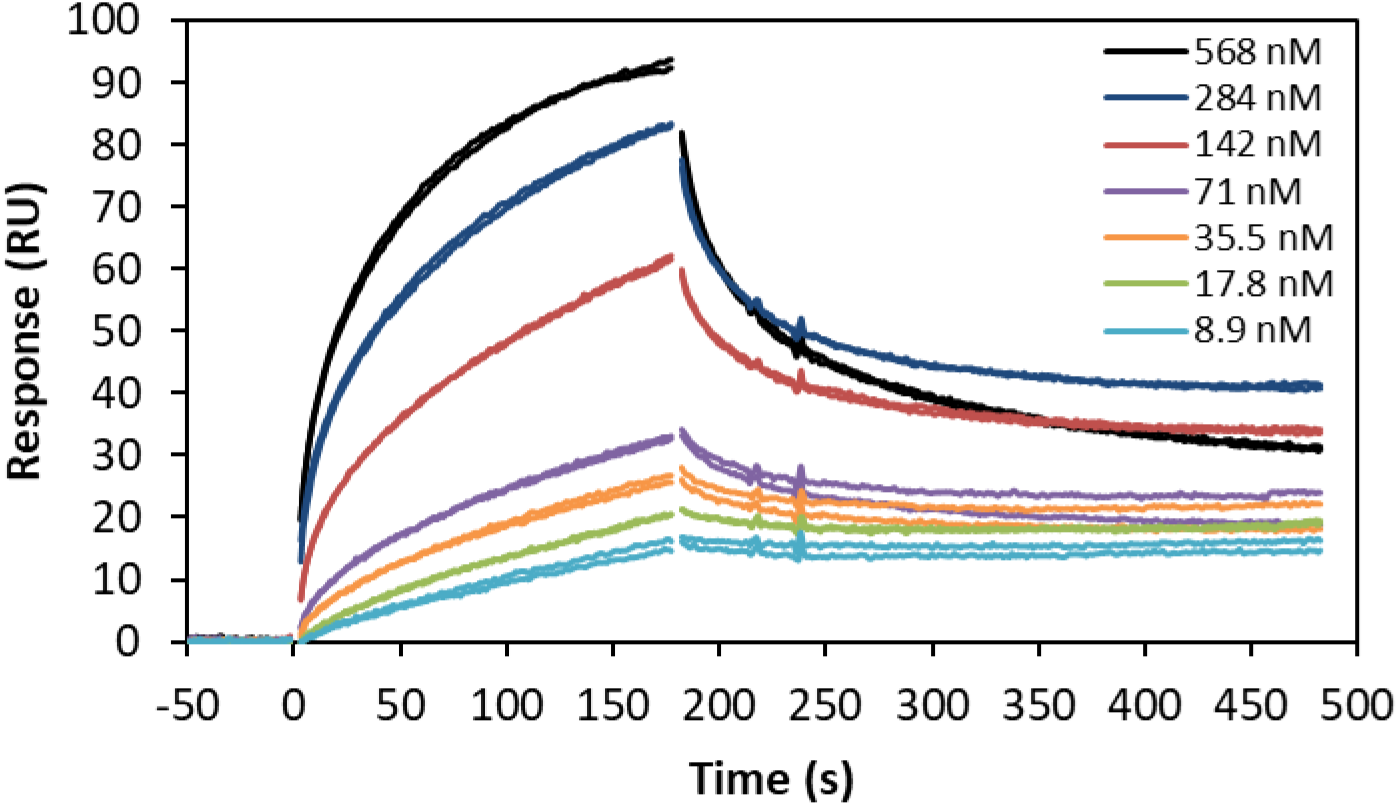
– Interaction assays between DsrC-trisulfide and DsrMKJOP complex. Sensorgrams of DsrC-trisulfide interaction with increasing concentrations of DsrMKJOP (8.9 nM to 568 nM) using SPR. Sensorgrams were measured in duplicate.

### The DsrC-trisulfide is reduced by DsrMKJOP

Given that menadiol can reduce the noncubane [4Fe-4S]^3+/2+^ catalytic center of DsrK, and that the DsrC-trisulfide interacts with DsrMKJOP complex, we tested whether the DsrC-trisulfide can serve as substrate and electron acceptor for the menadiol-reduced DsrMKJOP complex. Menadiol:DsrC-trisulfide oxidoreductase activity by the DsrMKJOP complex was thus measured. These experiments confirmed a fast oxidation of menadiol (measured by menadione formation at 270 nm) in the presence of the DsrC-trisulfide with a specific activity of 85.5 ± 5.4 mU/mg, assuming DsrMKJOP as a 5-subunit monomer (200 kDa) (**Figure 3A**). In control reactions without DsrMKJOP, menadiol or DsrC-trisulfide no activity was detected. In addition, sequential additions of DsrC-trisulfide, after the reaction has stopped, trigger a new phase of menadiol oxidation confirming it had stopped due to substrate depletion (**Figure 3B**).

**Figure 3.**
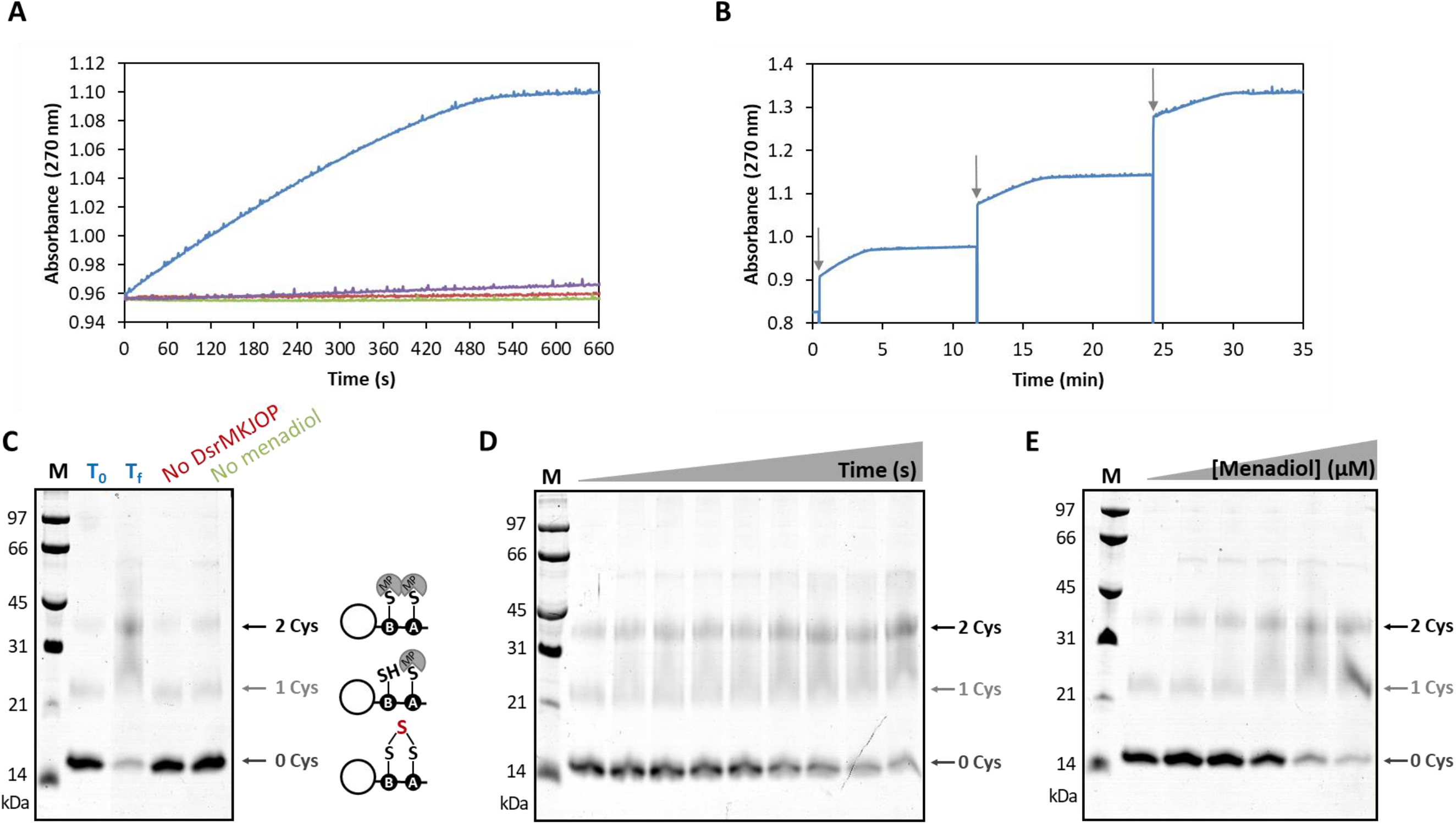
– Kinetic assays of the effect of DsrC-trisulfide on DsrMKJOP activity. (A) Menadiol:DsrC-trisulfide oxidoreductase activity catalyzed by *A. fulgidus* DsrMKJOP (blue), and in the absence of DsrC-trisulfide (purple), absence of DsrMKJOP (red), and absence of menadiol (green). (B) Menadiol:DsrC-trisulfide oxidoreductase activity catalyzed by *A. fulgidus* DsrMKJOP upon successive additions of DsrC-trisulfide (arrow indicates addition of 5 μM DsrC-trisulfide). (C and D) Gel-shift analysis of MalPEG-labeled DsrC from (A) and after reaction with DsrMKJOP and 0.5 mM menadiol over time (at 0, 1, 2, 3, 5, 10, 15, 20 and 30 min), respectively. MalPEG causes a shift of approximately 10 kDa per each labeled cysteine. (E) Gel-shift analysis of MalPEG-labeled DsrC with increasing concentrations of menadiol (0, 0.8, 4, 20, 100 and 500 μM), reacted for 30 mins. (M, molecular marker; T_0_, time zero; T_f_, time at the end of reaction).

The redox state of DsrC, before and after the reaction, was analyzed by a gel-shift assay using MalPEG, which selectively labels cysteine thiol groups (7, 37). In this assay, the DsrC-trisulfide presents no shift in the gel as it cannot bind MalPEG, whereas the reduced form binds one or two MalPEG molecules, as the last Cys_A_ is more accessible for labeling than Cys_B_ (Cys^114^ and Cys^103^ in *A. fulgidus*, respectively) (7, 37). At time zero (T_0_), most DsrC shows no shift, as expected for the trisulfide, but a small amount of reduced species is present, as revealed by labeling for one MalPEG, due to incomplete production of DsrC-trisulfide by DsrAB (**Figure 3C**). After the enzymatic reaction (T_f_), most DsrC is now in the reduced form, as revealed by the shift for one (Cys_A_) and two labeled cysteines (Cys_A_ and Cys_B_) (**Figure 3C**). In the control experiments, without DsrMKJOP or menadiol, DsrC remains in the oxidized DsrC-trisulfide state. Using the MalPEG assay to follow the reaction in time, the decrease in the DsrC-trisulfide band and increase in the band for reduced, doubly labeled DsrC, is clearly seen (**Figure 3D**). Increasing concentrations of menadiol for the same reaction time also increase the extent of reaction (**Figure 3E**). As a control, reduced DsrC was tested as a substrate for DsrMKJOP under the same conditions. In this case, no activity is observed (**Figure S2A**), as expected, and the MalPEG assay confirms the reduced state of DsrC in the beginning and end of the experiment (**Figure S2B).**

Finally, the redox state of DsrC cysteines was also characterized by peptide mass fingerprinting after protein digestion with Asp-N endoproteinase, which gives rise to a small peptide containing both Cys_A_ and Cys_B_, followed by alkylation with iodoacetamide (IA), which adds 57 Da per reduced cysteine (7). In reduced DsrC, the C-terminal peptide ^101^DA**C**RIAGLPKPTG**C**V^115^ has a mass of 1498.7 Da (7), and upon alkylation, a peptide with 1614.9 Da is detected (**Figure 4A**). In the DsrC-trisulfide formed by reaction with DsrAB, the C-terminal peptide shows a mass increase of 32 Da, 1530.7 Da, which remains unaltered by alkylation with IA (**Figure 4B**). A small amount of unreacted IA-labelled DsrC is still detected in this sample (1614.8 Da), as well as a species that does not react with IA (1498.8) and likely corresponds to DsrC with a disulfide bond between the two Cys. It should be noted that the peak intensities are not directly correlated with relative amounts, as they depend on the different ionization properties of each peptide. As expected, analysis of the DsrC peptide after the reaction with DsrMKJOP, reveals the disappearance of the DsrC-trisulfide peak at 1530.7 Da, and the increase of the 1614.8 Da peak characteristic of reduced DsrC, where two cysteines react with IA (**Figure 4C**).

**Figure 4.**
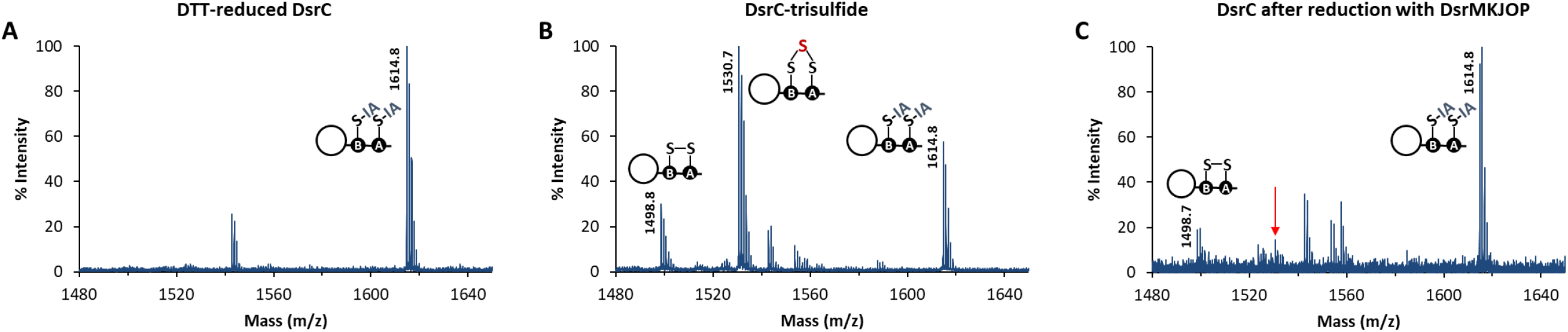
– Peptide mass fingerprinting spectra of DsrC C-terminal peptides after alkylation with IA. (A) DTT-reduced DsrC (before the reaction with *A. fulgidus* DsrAB and sulfite). (B) DsrC-trisulfide (produced in the *in vitro* enzymatic reaction with *A. fulgidus* DsrAB and sulfite). (C) DsrC recovered after the *in vitro* enzymatic reaction with *A. fulgidus* DsrMKJOP complex and menadiol. The C-terminal peptide of DsrC-trisulfide (1530.7 Da) is marked with an arrow in (C) showing its disappearance from (B) to (C). (IA, iodoacetamide; DTT, dithiothreitol).

### Sulfide is the product of DsrMKJOP catalysis

The products of DsrC-trisulfide reduction by the DsrMKJOP complex should be sulfide and reduced DsrC (**Figure 5A**). To quantify the sulfide released during DsrMKJOP catalysis, the reaction was performed with increasing concentrations of DsrC-trisulfide. These kinetic studies showed an increase in the length of the reaction with increasing substrate concentrations (**Figure 5B**), but not an increase in reaction rate. This indicates that the concentrations used are already saturating and that the Michaelis-Menten constant (*K*_M_) for the DsrC-trisulfide is probably in the nM range, in agreement with the measured K_D_. The reaction rates slow down as the substrate becomes exhausted. The product sulfide, quantified by HPLC as the monobromobimane fluorescent derivative, increases linearly with increasing DsrC-trisulfide concentrations (**Figure 5C**), although in sub-stoichiometric amounts (40 to 50%). This can be explained by two factors: the partial escape of hydrogen sulfide to the gas phase at pH 7.0, made worse by the high temperature required for the DsrMKJOP assay, and also by a degree of uncertainty associated with quantification of the DsrC-trisulfide. This quantification was performed indirectly using the BCA (for protein quantification) and DTNB (for thiol quantification) assays, by the amount of reduced DsrC remaining in samples of the DsrC-trisulfide. However, we observed by mass spectrometry (MS) that upon production of the DsrC-trisulfide some part of DsrC is in the disulfide state and is not alkylated by IA (peak 1498.8 Da in **Figure 4B and 4C**) and thus is also not quantified by DTNB derivatization, generating an uncertainty in the actual level of DsrC-trisulfide present in the samples. Nevertheless, taken globally these results show clearly that DsrMKJOP can catalyze full reduction of the DsrC-trisulfide to sulfide using menadiol, finally clarifying the functional role of this respiratory complex in dissimilatory sulfite reduction.

**Figure 5.**
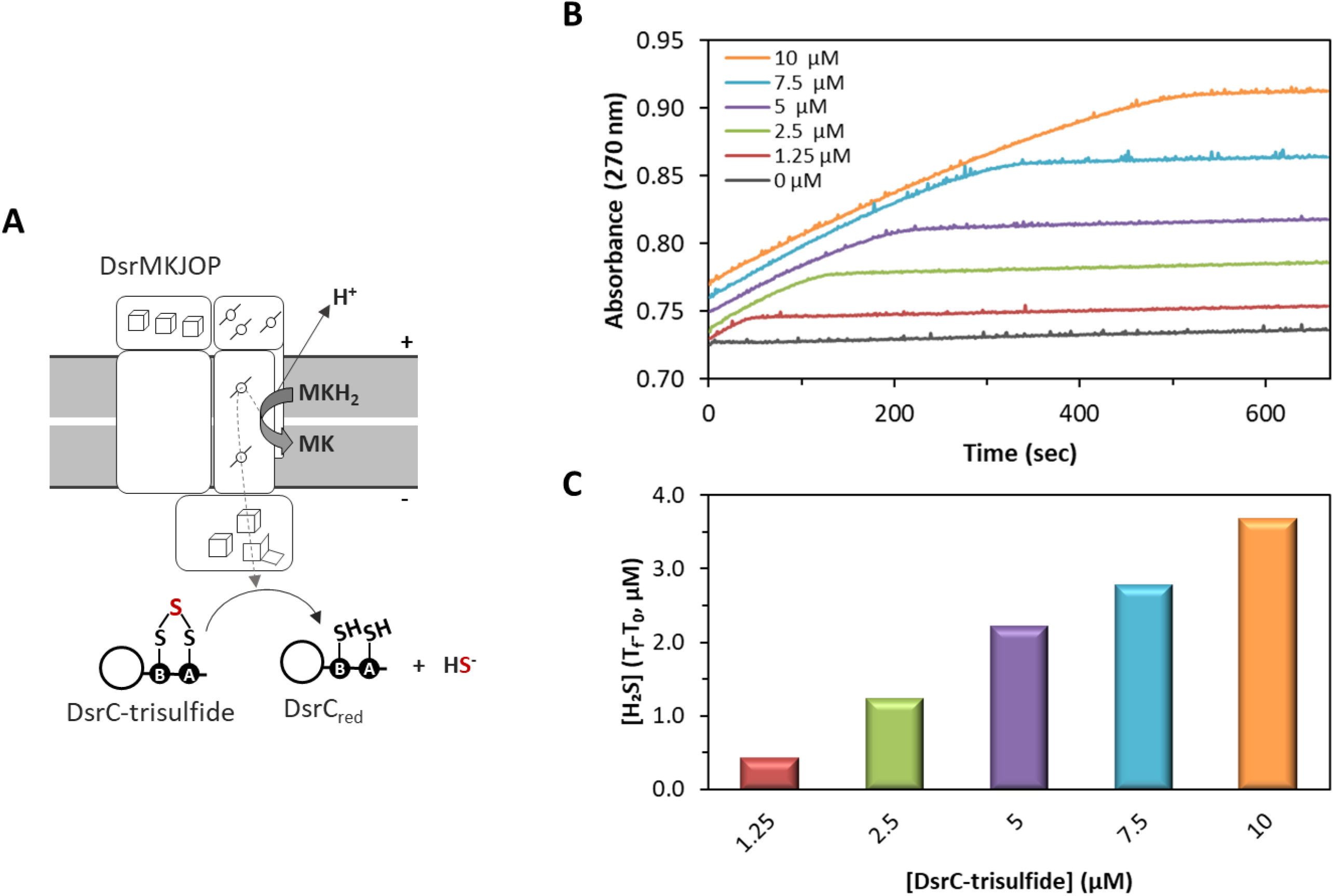
– Reduction of DsrC-trisulfide by DsrMKJOP and sulfide production. (A) Schematic representation of the reaction catalyzed by DsrMKJOP. (B) Menadiol:DsrC-trisulfide oxidoreductase activity catalyzed by *A. fulgidus* DsrMKJOP with increasing concentrations of DsrC-trisulfide (0 μM to 10 μM). (C) Hydrogen sulfide produced in the reactions shown in (B). H_2_S was measured before the addition of DsrC-trisulfide (T_0_) and 30 min after the start of the enzymatic reaction (T_f_). The control in the absence of DsrC-trisulfide was subtracted from all points. Results from a representative experiment are shown.

### Insights into the catalytic mechanism of DsrMKJOP

A catalytic mechanism for heterodisulfide reduction by Hdrs has been proposed based on the structure of HdrABC with substrate bound, where the CoM-S-S-CoB heterodisulfide binds between the two noncubane [4Fe-4S]^3+/2+^ clusters and is homolytically cleaved (30). The thyil radicals formed may be stabilized by coordination with the reduced [4Fe-4S]^2+^ clusters, leading to the binding of the CoM and CoB sulfurs to the two [4Fe-4S]^3+^ clusters, resulting in the “open” conformation of the clusters (31). Two consecutive single-electron reduction steps release CoB-SH and CoM-SH (30, 31). However, the catalytic mechanism of the DsrMKJOP complex is likely different since it coordinates a single noncubane [4Fe-4S]^3+/2+^ cluster in DsrK instead of the two found in HdrD and HdrB (**Figure S3**). Interestingly, in many DsrK proteins from different sulfate reducers the last cysteine of the CCG motif for binding this cluster is replaced by a conserved aspartate (**Figure S4**). In *A. fulgidus*, the DsrK protein of the DsrMKJOP complex (AF0502) and another homolog (AF0543) have this aspartate, whereas two other DsrK homologs (AF0544 and AF0547), encoded in *dsrMK* arrangements, have a cysteine, suggesting that both residues may be functional. In addition, one of the conserved cysteines from the missing noncubane cluster (Cys^329^ in *A. fulgidus* numbering) is strictly conserved in DsrK proteins and is likely involved in the catalytic mechanism of DsrC-trisulfide reduction. This cysteine is part of a conserved GECGH sequence motif that is highly conserved in DsrK from several organisms (**Figure S4**) and is one of the cysteines from the second CCG signature motif in HdrD and HdrB (**Figure S3**). A similar motif (ECGLH), present in APS and PAPS reductases, includes a redox-active catalytic cysteine (44). Other proteins from sulfate reducers, homologous to DsrK and HdrD, such as HmcF from the HmcABCDEF complex (19, 24, 45) and TmcB from the TmcABCD complex (46) also contain only one CCG motif and this conserved cysteine in a T/GECGH motif (**Figure S3**), suggesting that this cysteine is probably essential for disulfide reduction catalysis. The Hmc and Tmc complexes are both transmembrane complexes with a similar architecture to the DsrMKJOP complex, but they are not present in *A. fulgidus* (19). Interestingly, the characteristic EPR signal of the noncubane [4Fe-4S]^3+^ cluster was initially reported for the methanogenic HdrB and HdrD only in the presence of HS-CoM or HS-CoB (41–43), which bind to the cluster, whereas for DsrK and TmcB proteins this signal is observed in the as-isolated protein, in the absence of any added substrate (11, 16, 46). This could suggest that this conserved cysteine might be binding the cluster and generating the open *S* = ½ conformation. However, for the recombinant HdrB protein produced in *E. coli* the signal for the noncubane cluster was visible already in the absence of substrate (23).

In methanogens, the redox potential of the CoM-S-S-CoB heterodisulfide has been determined to be *E*^0^’=-143 ± 10 mV vs. SHE (47), which is more positive than that of typical disulfides such as glutathione and cysteine. It is also more positive than the potential of methanophenazine (*E*^0^’=-165 ± 6 mV), the membrane-associated quinone-like redox carrier that is the electron donor to HdrED (47). The redox potential of the DsrC-trisulfide is unknown, so it is not clear whether its reduction by menaquinol (*E*^0^’ ∼ - 70 mV) is thermodynamically favorable. Protein disulfide bonds are known to span a wide range of potentials. For example, *A. fulgidus* proteins from the thioredoxin superfamily have potentials ranging from -32 mV to -309 mV (pH 7.0) (48). A value of -346 mV (pH 7.0, vs. SHE) has been reported for several organic water-soluble trisulfides (49), but we know of no reports for redox potentials of protein-bound trisulfides, so it is very difficult to estimate the redox potential of the DsrC trisulfide. We tested the chemical reduction of this species by different chemical reducing agents, and surprisingly none of them, not even those with very low redox potential, could reduce the DsrC-trisulfide as analyzed with the MalPEG shift assay (**Figure S5**). Moreover, MS confirmed the intact state of the DsrC-trisulfide after reduction with dithionite, TCEP and DTT (**Table S1**). This unexpected result suggests that the DsrC-trisulfide can only be reduced by the DsrMKJOP complex. The fact that it cannot be chemically reduced could suggest that it has a low redox potential, which could increase upon interaction with the DsrMKJOP complex, but it is also possible that the reduction of the trisulfide is not thermodynamically, but kinetically prevented, i.e. that the reaction has a high activation energy and occurs only when it is enzymatically catalyzed. This sort of argument has been put forward by Flohé and coworkers in discussing the glutathione redox potential and its physiological significance (50, 51). Such kinetic impediment for reduction of the DsrC-trisulfide, which may be based on steric inaccessibility of the trisulfide, makes sense considering that this species is likely to face a considerable intracellular sulfide concentration.

Nevertheless, the full reduction of the DsrC-trisulfide, as observed by MS, and the increasing production of sulfide upon reaction of DsrMKJOP and menadiol with increasing concentrations of DsrC-trisulfide confirm that its reduction by this menaquinol analog does occur. This reaction could, in principle, only involve the DsrMK subunits in analogy to HdrED, with DsrM as the menaquinol-oxidizing subunit that transfers electrons to DsrK for trisulfide reduction, and in fact, in many organisms only these two subunits are present (2, 19). This leaves the function of the DsrJOP subunits as an enigma.

It has been suggested that the DsrJ cytochrome could react with a periplasmic sulfur substrate (11, 16, 25), due to the presence of the unusual His/Cys heme coordination that is also present in SoxXA involved in thiosulfate oxidation (52) or in TsdA thiosulfate dehydrogenase (27). We tested both thiosulfate and sulfite as electron donors to the complex and could not detect any activity with menadione and/or DsrC-trisulfide as electron acceptors, and sulfite actually decreased the rate of DsrC-trisulfide reduction by DsrMKJOP (**Figure S6**). Since the DsrJOP module is not essential for some organisms, a hypothesis is that its role is to increase the energetic efficiency of the DsrMKJOP complex vs. the DsrMK version. We have previously suggested that the DsrMK and DsrJOP modules could be involved in some form of proton-translocating quinone cycling, as present in the *bc*_1_ complex, possibly involving quinone-based electron bifurcation (10). Alternatively, the DsrJOP module may allow for proton pumping upon DsrC-trisulfide reduction. Notably, the DsrOP proteins form a quinone-interacting redox module that is widespread in bacterial oxidoreductases (24, 53), and some proteins of the DsrP family may be involved in proton pumping, such as the alternative complex III (54, 55) and the reductive dehalogenase complex (56).

In conclusion, our results are a major step forward in the molecular understanding of dissimilatory sulfur metabolism by identifying the function of the DsrMKJOP complex, showing that it interacts with the DsrC-trisulfide and acts as a menadiol:DsrC-trisulfide oxidoreductase that produces sulfide. The architecture of the DsrMKJOP complex indicates that it is quite an elaborate machine and further studies will be essential to understand its molecular mechanism and how it is coupled to energy conservation.

## MATERIALS AND METHODS

### Purification of *A. fulgidus* DsrMKJOP

*A. fulgidus* VC-16 (DSM 4304) was grown in sulfate-thiosulfate-lactate medium in 300 L fermenter at 83 °C, as previously described (32), without addition of dithionite. Frozen cells were resuspended in 50 mM potassium phosphate buffer (KPi) pH 7.0, 10% (v/v) glycerol, homogenized and disrupted in an APV Model 2000 Homogenizer at 80 MPa in the presence of DNase (Sigma-Aldrich) and cOmplete^TM^ protease inhibitor cocktail (Roche). The cell lysate was centrifuged at 7930 x *g* for 20 min at 4 °C, followed by ultracentrifugation at 138000 x *g* for 2 h at 4 °C. The supernatant (soluble extract) was used to isolate *A. fulgidus* DsrAB (see below). The pellet (membrane extract) was solubilized with gentle stirring overnight at 4 °C in 50 mM potassium phosphate buffer pH 7.0, 10% (v/v) glycerol, and 2% (w/v) n-dodecyl β-D-maltoside (DDM, Glycon Biochemicals GmbH). The solubilized membrane proteins were separated by ultracentrifugation at 138000 x *g* for 2 h at 4 °C, and the membrane pellet was used for a second solubilization for 5 h with gentle stirring at 4 °C, followed by a third ultracentrifugation step under the same conditions. The pooled solubilized membrane proteins were loaded into a Q-Sepharose high performance column (2.6 x 10.0 cm, Amersham Pharmacia Biotech) equilibrated with buffer A (50 mM KPi pH 7.0, 10% (v/v) glycerol, 0.1% (w/v) DDM, and a cOmplete^TM^ protease inhibitor tablet/L). A stepwise gradient of increasing concentrations of NaCl (from 0 to 1 M NaCl) was applied and the fractions eluted at 0.45 M and 0.5 M NaCl were pooled, concentrated and the ionic strength of the solution was lowered by dilution with buffer A and ultrafiltration (50 kDa cutoff, Amicon, Millipore). The concentrate was applied to a Resource Q column (1.6 x 3.0 cm, Amersham Pharmacia Biotech) equilibrated with buffer A. A stepwise gradient of increasing concentrations of NaCl (from 0 to 1 M NaCl) was performed and the fractions eluted at 0.3 M and 0.35 M NaCl were pooled, concentrated and the ionic strength lowered by dilution and concentration steps as above. All the chromatographic steps were performed at 4 °C, at air, and monitored by UV-visible spectroscopy. The purity of DsrMKJOP was analyzed by a 10% Tricine-SDS-PAGE gel and a Blue Native (BN)-PAGE gel, followed by Coomassie Blue staining and heme-staining (33), and the protein concentration was determined using the previously determined absorption coefficient (ε) of 132.4 mM⁻¹.cm⁻¹ at 555 nm in the reduced state (34).

### UV-visible Spectroscopy

UV–visible spectra were acquired in a Shimadzu UV-1800 spectrophotometer inside a Coy anaerobic chamber (98 % N_2_, 2 % H_2_). Spectra were obtained at 60 °C, using as-purified DsrMKJOP, prepared in 20 mM Tris-HCl pH 7.5, then incubated with menadiol and an excess of sodium dithionite to allow full reduction. Menadiol was prepared by reduction of menadione with sodium dithionite (35), stored in dry state at -20 °C under anaerobic conditions, and solubilized in ethanol prior to use. A control spectrum of the as-purified DsrMKJOP complex was obtained by addition of the same volume of ethanol as for menadiol incubation.

### Electron Paramagnetic Resonance Spectroscopy

X-band electron paramagnetic resonance (EPR) spectra were obtained on a Bruker EMX spectrometer (Billerica, MA) equipped with an Oxford Instruments ESR-900 continuous flow helium cryostat (Abingdon, UK). A menadiol-treated sample of the DsrMKJOP complex, in 20 mM Tris-HCl pH 7.5 at a concentration of 25 μM, was prepared inside a Coy anaerobic chamber (98 % N_2_, 2 % H_2_) by incubation for 2h at 60 °C with a solution of menadiol in ethanol in a 1:500 ratio. As control an as-purified DsrMKJOP sample was also incubated for 2 h with the same volume of ethanol as was used in the menadiol experiment. Samples were transferred to EPR tubes within the anaerobic chamber, capped, and immediately frozen in liquid nitrogen upon removal from the chamber. All spectra were recorded using a microwave frequency of 9.38 GHz, a field modulation frequency of 100 kHz, a microwave power of 20.1 mW, a modulation amplitude of 1.0 mT, and a temperature of 23 K.

### Purification of *A. fulgidus* DsrAB

The purification protocol of the soluble extract from *A. fulgidus* cells followed two consecutive ion exchange chromatography steps at 4 °C under air, as described in detail in Santos *et al.* (7). The purity of DsrAB was analyzed in a 10% Tricine-SDS-PAGE gel, and the protein concentration was determined by UV-visible spectroscopy using an ε of 60 mM⁻¹.cm⁻¹ at 593 nm determined previously (36). The purified DsrAB was stored under anaerobic conditions to prevent prolonged contact with O_2_, which leads to gradual activity loss.

### Purification of Recombinant *A. fulgidus* DsrC

*A. fulgidus* DsrC with the two structural Cys (Cys^77^ and Cys^85^) modified to Ala was expressed in *Escherichia coli* and purified by an affinity exchange chromatography step on a HiTrap^TM^ immobilized metal-ion affinity chromatography (IMAC) high performance column (GE Healthcare), as described in detail in Ferreira *et al.* (9). The purity of DsrC was analyzed on a 10% Tricine-SDS-PAGE gel, and the protein concentration was determined by bicinchoninic acid (BCA) assay (Merck Millipore) and by UV-visible spectroscopy using an ε of 24 mM⁻¹·cm⁻¹ at 280 nm. Before its use in activity assays, DsrC was treated with 5 mM dithiothreitol (DTT, Sigma-Aldrich) during 30 min at 37 °C, to ensure its Cys were in the reduced state, and excess reductant was removed with a HiTrap^TM^ Desalting column (GE Healthcare) (7). Reduced DsrC was concentrated, if necessary, by ultrafiltration (10 kDa cutoff, Amicon Millipore) and kept under anaerobic conditions until use. The redox state of the DsrC (5 μg) was analyzed by a gel-shift assay using MalPEG (methoxy-polyethylene glycol maleimide, Fluka), as described in Venceslau *et al.* (37).

### Production and Analysis of DsrC-trisulfide

The enzymatic production of DsrC-trisulfide was performed inside a Coy anaerobic chamber (98% N_2_, 2% H_2_) at 60 °C using Zn-reduced methyl viologen (Sigma-Aldrich) as the electron donor (7). The activity assays were run in 50 mM KPi pH 7.0, 1 mM Zn-reduced methyl viologen (MV^+^), 200 nM of DsrAB, 0.5 mM sodium sulfite (Sigma-Aldrich), with a range from 10 μM to 100 μM of reduced DsrC. After a 5 min incubation of DsrAB in the reaction buffer, followed by a 1 min incubation with DsrC, the reaction was started by addition of sodium sulfite. The oxidation of MV⁺ was monitored at 732 nm (ε = 3.15 mM⁻¹·cm⁻¹) in a spectrophotometer. A prolonged incubation time (from 15 to 60 min depending on the DsrC-trisulfide concentration) was used to try to drive the enzymatic reaction to completion and have DsrC as much as possible in the trisulfide form. The enzymatic solution was diluted and concentrated by ultrafiltration (10 kDa cutoff, Amicon Millipore) to remove the methyl viologen. The redox state of DsrC was monitored with the MalPEG gel-shift assay as previously described (37).

### Determination of DsrC Thiol Content by DTNB

DsrC in different redox states was subject to a DTNB (5,5-dithio-bis-(2-nitrobenzoic acid) – Ellman’s reagent) assay to quantify the sulfhydryl groups present. DsrC (0.8 – 3 nmol) was incubated in 100 mM KPi pH 8.0, 1 mM EDTA, and 0.2 mM DTNB (Sigma-Aldrich) for 15 min at room temperature. The production of 2-nitro-5-thiobenzoate anion (TNB^2-^) was quantified by measuring the absorbance at 412 nm using the ε of 14150 M⁻¹·cm⁻¹ (38).

### Surface Plasmon Resonance Analysis

The surface plasmon resonance (SPR) experiments were performed at 25 °C on a BIAcore 2000 instrument (GE Healthcare). DsrC-trisulfide was immobilized on a NTA sensor chip (GE Healthcare) by the His-tag tail present at the N terminus. All the assays were performed with 10 mM HEPES pH 7.4, 150 mM NaCl, 50 μM EDTA, and 0.005% (v/v) Tween 20 running buffer. DsrC-trisulfide (45 nM) was bound to a previously activated flow cell at a flow rate of 5 μL/min in the presence of 500 μM NiCl_2_ in running buffer in order to have an immobilization of around 350 resonance units (RU). Another activated flow cell was similarly treated with running buffer in the absence of DsrC-trisulfide (control cell). Interaction experiments with DsrMKJOP complex were performed with a 3 min injection of increasing concentrations of DsrMKJOP complex (range from 8.9 nM to 568 nM) at a flow rate of 30 μL/min. At the end of sample injection, the NTA sensor chip was regenerated with 350 mM EDTA in running buffer for 1 min at a flow rate of 10 μL/min before a new cycle of surface activation and immobilization. Sensorgrams were obtained by subtracting the unspecific binding to the control flow cell in order to remove buffer artifacts and normalizing to the baseline injection timepoint. The equilibrium dissociation constant (K_D_) was determined from duplicate experiments and according to RU_steady-state_ = (RU_max_ x [analyte])/K_D_ + [analyte]).

### DsrMKJOP Activity Assays

Menadiol:DsrC-trisulfide oxidoreductase activity assays by DsrMKJOP were performed inside a Coy anaerobic chamber (98% N_2_, 2% H_2_) at 60 °C using menadiol as electron donor. The activity assays were run in 50 mM KPi pH 7.0, 0.5 mM menadiol, and 100 nM DsrMKJOP in the presence of DsrC-trisulfide (10 μM). The reaction was started by addition of DsrC-trisulfide after a 5 min incubation of DsrMKJOP in the reaction buffer. The oxidation of menadiol was monitored at 270 nm (using ε_DMN_ = 16 mM⁻¹·cm⁻¹ (39)) in a spectrophotometer. For peptide mass fingerprinting of DsrC, the enzymatic reactions were concentrated and dialyzed by ultrafiltration (10 kDa cutoff, Amicon Millipore). To monitor the redox state of DsrC cysteines, the MalPEG gel-shift assay was used (37).

### Peptide Mass Fingerprinting

The redox state of DsrC cysteines was analyzed by peptide mass fingerprinting using a matrix-assisted laser desorption/ionization-time-of-flight (MALDI-TOF/TOF) mass spectrometer, as described in Santos *et al.* (7). Alkylation was performed inside an anaerobic chamber by incubation of DsrC with 50 mM iodoacetamide (IA, Sigma-Aldrich) in 50 mM ammonium bicarbonate pH 7.8 during 30 min at 55 °C and run on a 10% Tricine-SDS-PAGE gel (15 μg) under non-reducing conditions. The protein bands were excised, destained and digested with Asp-N (Roche, 20 ng/μL) overnight at 37 °C. This endoproteinase generates a peptide comprising both conserved C-terminal cysteine residues, ^101^DA**C**RIAGLPKPTG**C**V^115^, with predicted mass of 1500.8 Daltons (Da). The digested peptides were desalted and concentrated using POROS C18 (Empore, 3M) and eluted directly onto the MALDI plate using 1 μL of 5 mg/mL alpha-cyano-4-hydroxycinnamic acid (CHCA, Sigma-Aldrich) in 50% (v/v) acetonitrile and 5% (v/v) formic acid.

The data were acquired in positive reflector MS mode using a 5800 MALDI-TOF/TOF (AB Sciex) mass spectrometer and TOF/TOF Series Explorer Software v.4.1.0 (Applied Biosystems). External calibration was performed using CalMix5 (Protea). Mass Spectrometry (MS) data were analyzed using Data Explorer Software v.4.11 (AB Sciex) and obtained at the Mass Spectrometry Unit (UniMS), ITQB/iBET, Oeiras, Portugal.

### Determination of Hydrogen Sulfide by HPLC

Hydrogen sulfide was quantified using monobromobimane (mBBr) derivatization (40). mBBr (Thermo Fisher Scientific) was diluted in HPLC ultrapure acetonitrile to a final concentration of 50 mM. For derivatization, 10 μL of sample from the enzymatic reaction were reacted with 3 μL of mBBr in 50 mM EPPS (4-(2-hydroxyethyl)piperazine-1-propanesulfonic acid) pH 8.0 and 5 mM DTPA (diethylenetriaminepentaacetic acid) in a final volume of 86 μL, inside an anaerobic chamber. The reactions were incubated in the dark at room temperature for 10 min, after which 4 μL of 5 M methanesulfonic acid was added to stop the reaction. Derivatized hydrogen sulfide (sulfide dibimane) was measured by high performance liquid chromatography (HPLC, Waters Alliance 2695) with a fluorescence detector (Waters 2475) with an excitation/emission wavelength of 380 and 480 nm, respectively, using a reverse phase Ultrasphere® ODS column (4.6 x 250 mm, 5 μm, Hichrom) run at a flow rate of 1.2 mL·min⁻¹ and 35 °C, as described in detail in Santos *et al.* (7). The retention time was 15.5 min for sulfide dibimane. A calibration curve was performed with freshly prepared solutions of sodium sulfide nonahydrate (Sigma-Aldrich) in 50 mM KPi pH 7.0, reacted with excess mBBr.

## Supporting information

Supplementary information DsrMKJOP

## ACKNOWLEDGEMENTS

We thank João Carita and Cristina Leitão from ITQB NOVA for the growth of *A. fulgidus* and HPLC analysis, respectively. This work was funded by the Fundação para a Ciência e Tecnologia (Portugal) through Fellowships PD/BD/135488/2018 and COVID/BD/152504/2022 (A.C.C.B.), Grants PTDC/BIA-MIC/6512/2014 and PTDC/BIA-BQM/29118/2017, Research Unit Molecular, Structural and Cellular Microbiology (MOSTMICRO-ITQB) (UIDB/04612/2020 and UIDP/04612/2020), and Associate Laboratory Life Sciences for a Healthy and Sustainable Future (LS4FUTURE) (LA/P/0087/2020).

