## Supplementary information DsrMKJOP for "DsrMKJOP is the terminal reductase complex in anaerobic sulfate respiration"

for

**Table S1 – Peptide mass fingerprinting data of the DsrC C-terminal peptide, before and after alkylation with IA.** The analysis was performed on DTT-reduced DsrC and DsrC-trisulfide as controls, and after the attempted reduction of the DsrC-trisulfide (0.13 mM) for 2 h at 25 °C in the presence of 2 mM of each reducing agent (sodium dithionite, TCEP and DTT). (IA, iodoacetamide; TCEP, Tris(2-carboxyethyl)phosphine; DTT, dithiothreitol).

| | **-IA**<br>**Mass (Da)** | **+IA**<br>**Mass (Da)** | **Notes** |
|---|---|---|---|
| **DTT-reduced DsrC (Control)** | **1498.0** | **1614.7** | 2 Cys available |
| **DsrC-trisulfide (Control)** | **1530.4**<br>(minor 1498.4) | **1529.6**<br>(No 1614.7 species) | DsrC trisulfide present |
| **DsrC-trisulfide + Dithionite** | **1529.5**<br>(minor 1497.5) | **1529.4**<br>(No 1614.7 species) | DsrC trisulfide present |
| **DsrC-trisulfide + TCEP** | **1531.4**<br>(minor 1497.5) | **1530.1**<br>(No 1614.7 species) | DsrC trisulfide present |
| **DsrC-trisulfide + DTT** | **1531.4**<br>(minor 1499.2) | **1531.1**<br>(No 1614.7 species) | DsrC trisulfide present |

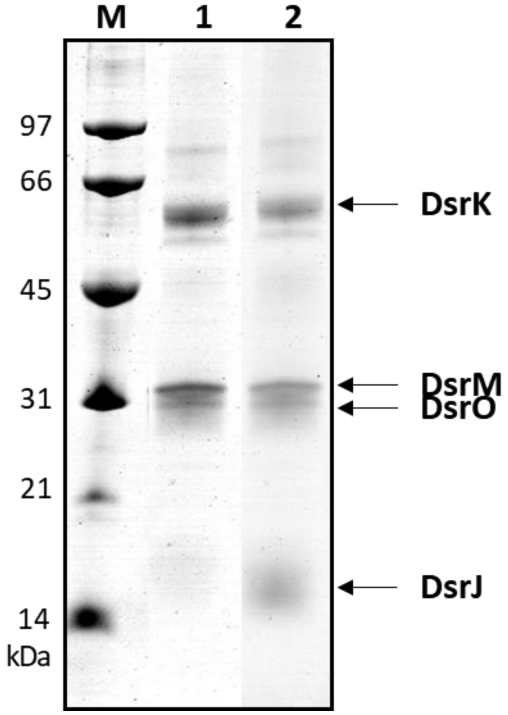


**Figure S1 – Tricine-SDS-PAGE gel of as-purified *A. fulgidus* DsrMKJOP complex.** (M) Molecular marker, (1) Coomassie Blue stained *A. fulgidus* DsrMKJOP complex (15 µg), followed by heme staining of lane 1 (2). The complex was denaturated for 15 min at 37 °C in SDS loading buffer under non-reducing conditions.

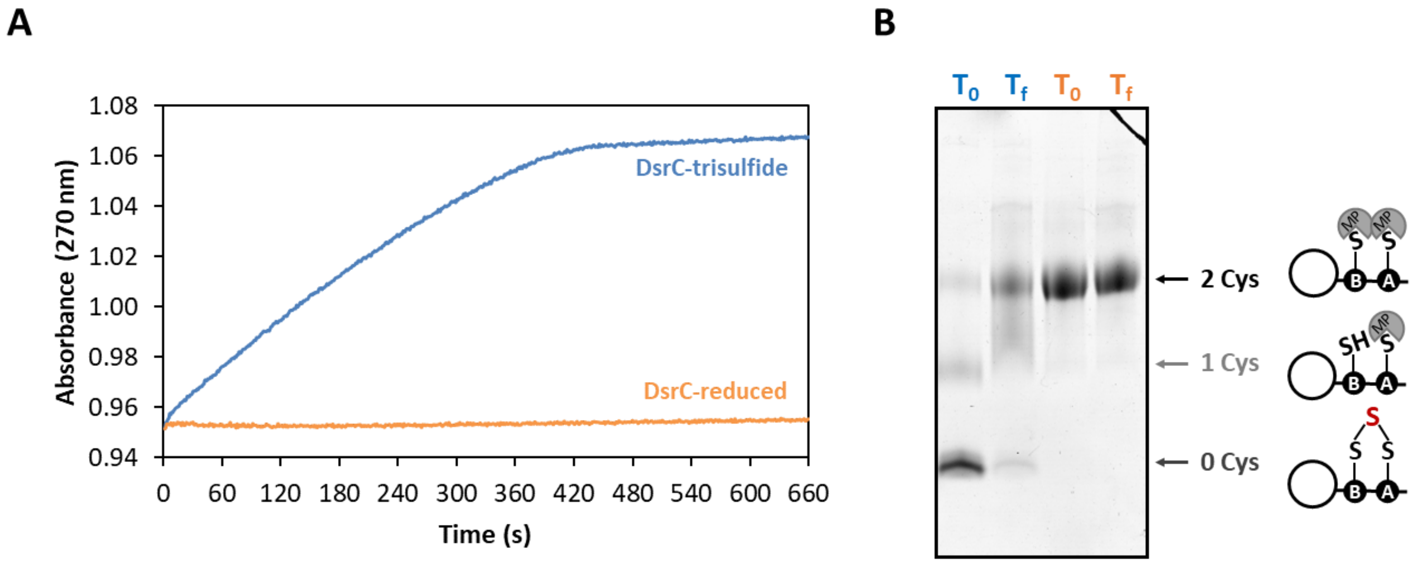


**Figure S2 – Control reaction between DsrMKJOP and reduced DsrC.** (A) Menadiol:DsrC-trisulfide oxidoreductase activity catalyzed by *A. fulgidus* DsrMKJOP (blue) versus reaction in the presence of reduced DsrC (orange). (B) Gel-shift analysis of MalPEG-labelled DsrC from (A).

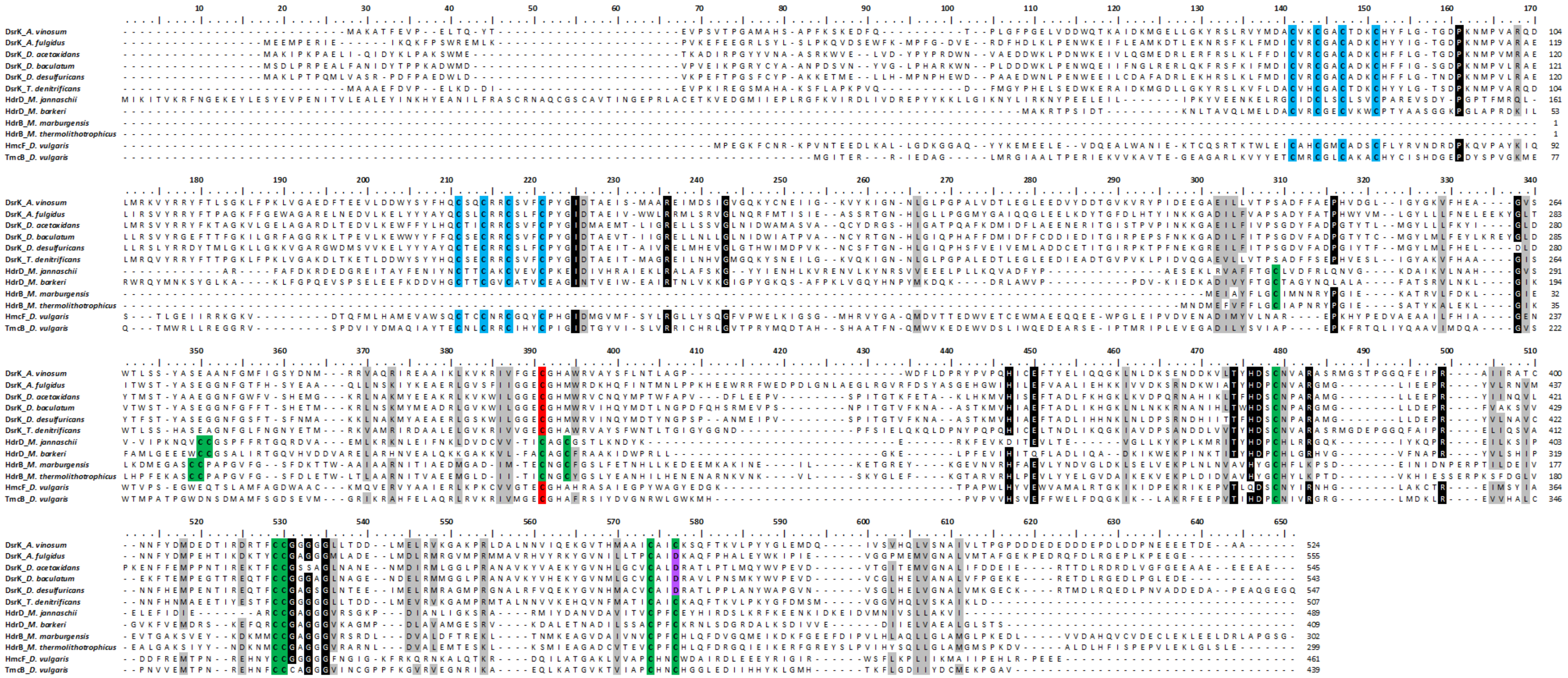


**Figure S3 – DsrK, HdrD, HdrB, HmcF and TmcB multiple sequence alignment.** Sequences were downloaded from the Joint Genome Institute (https://img.jgi.doe.gov/) and aligned with Clustal Omega (https://www.ebi.ac.uk/Tools/msa/clustalo/). Cysteines from the canonical $[4Fe\text{-}4S]^{2+/1+}$ clusters are highlighted in blue, and from the noncubane $[4Fe\text{-}4S]^{3+/2+}$ clusters in green. In DsrK of some organisms the last cysteine from the noncubane $[4Fe\text{-}4S]^{3+/2+}$ cluster is replaced by an aspartate, which is marked in purple. The conserved cysteine residue from the GECGH sequence motif is marked in red. Other conserved residues are marked in black (threshold 80% identity). Proteins are from *Allochromatium vinosum*, *Archaeoglobus fulgidus*, *Desulfobacca acetoxidans, Desulfomicrobium baculatum*, *Desulfovibrio desulfuricans, Thiobacillus denitrificans*, *Methanocaldococcus jannaschii*, *Methanosarcina barkeri*, *Methanothermobacter marburgensis*, *Methanothermococcus thermolithotrophicus* and *Desulfovibrio vulgaris* Hildenborough.

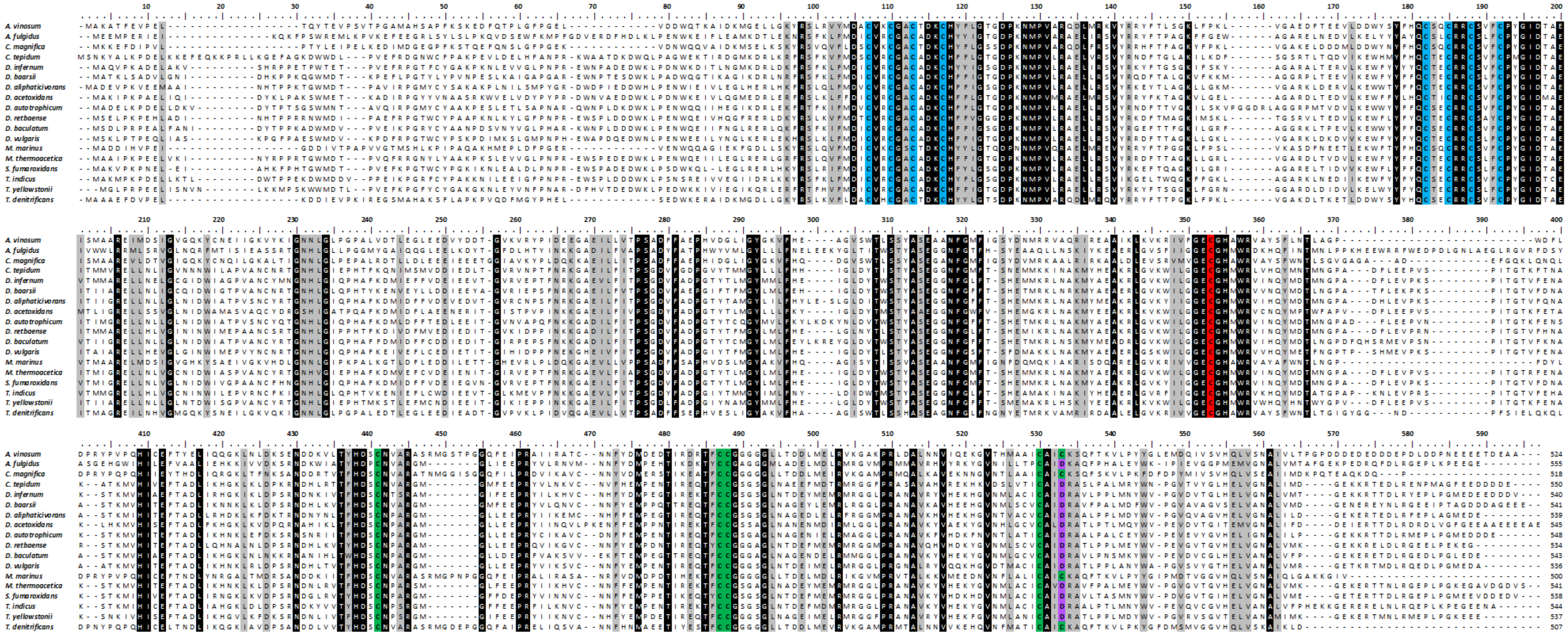

**Figure S4 – DsrK multiple sequence alignment.** Sequences were downloaded from the Joint Genome Institute (https://img.jgi.doe.gov/) and aligned with Clustal Omega (https://www.ebi.ac.uk/Tools/msa/clustalo/). Cysteines from the canonical $[4Fe\text{-}4S]^{2+/1+}$ clusters are marked in blue, and from the noncubane $[4Fe\text{-}4S]^{3+/2+}$ cluster in green. In DsrK of some organisms the last cysteine from the noncubane $[4Fe\text{-}4S]^{3+/2+}$ cluster is replaced by an aspartate, which is marked in purple. The conserved cysteine residue from the GECGH sequence motif is marked in red. Other conserved residues are marked in black (threshold 100% identity). Proteins are from *Allochromatium vinosum*, *Archaeoglobus fulgidus*, *Candidatus Ruthia magnifica*, *Chlorobaculum tepidum*, *Desulfacinum infernum*, *Desulfarculus baarsii*, *Desulfatibacillum aliphaticivorans*, *Desulfobacca acetoxidans*, *Desulfobacterium autotrophicum*, *Desulfohalobium retbaense*, *Desulfomicrobium baculatum*, *Desulfovibrio vulgaris* Hildenborough, *Magnetococcus marinus*, *Moorella thermoacetica*, *Syntrophobacter fumaroxidans*, *Thermodesulfatator indicus*, *Thermodesulfovibrio yellowstonii*, and *Thiobacillus denitrificans*.

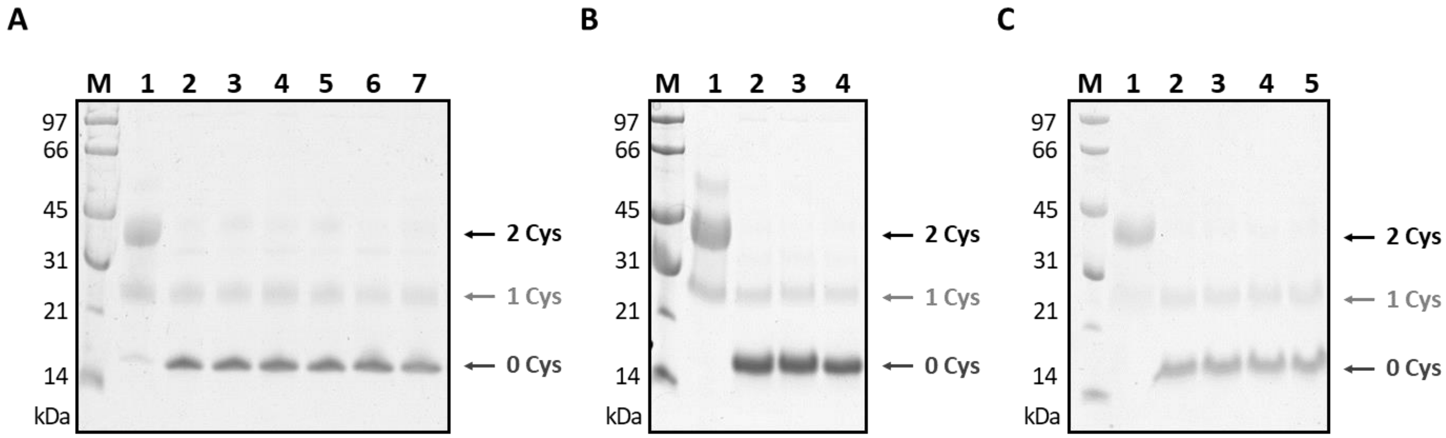


**Figure S5 – Effect of different chemical reducing agents on DsrC-trisulfide by gel-shift analysis using MalPEG.** In the three gels, lanes 1 and 2 are control samples: reduced DsrC (1) and DsrC-trisulfide (2). (A) Samples after DsrC-trisulfide reduction for 2 h at 25 °C in the presence of 2 mM of each reducing agent, namely sodium dithionite (3), 2-mercaptoethanol (4), tris(2-carboxyethyl)phosphine (5), dithiothreitol (6) and sodium sulfide nonahydrate (7). (B) Reduction of DsrC-trisulfide for 2 h at 25 °C in the presence of 2 mM of each reducing agent, namely sodium borohydride [1] (3) and titanium(III) citrate [2] (4). (C) Reduction of DsrC-trisulfide for 1 h at 50 °C in the presence of 20 mM (3) or 100 mM of sodium borohydride (4) or with zinc granules (5). All assays were performed under anaerobic conditions.

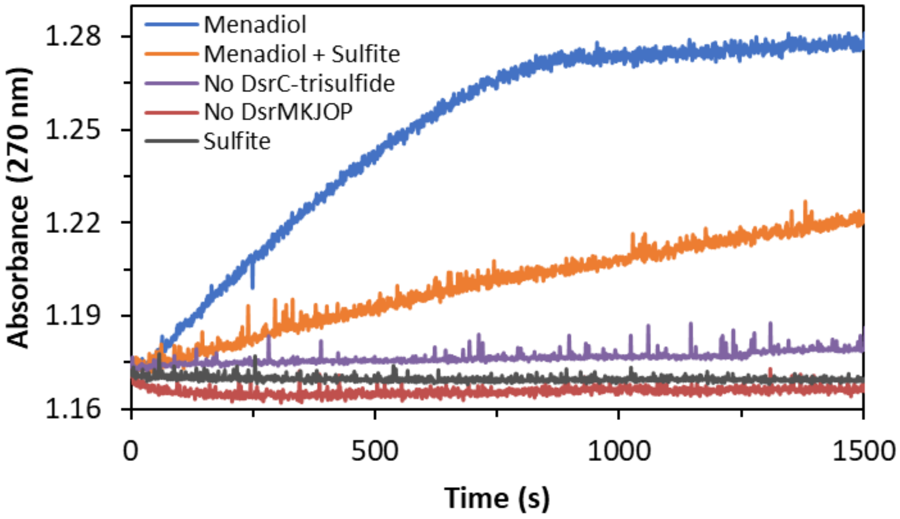


**Figure S6 –** Kinetic assays of the effect of DsrC-trisulfide on DsrMKJOP activity. Menadiol:DsrC-trisulfide oxidoreductase activity catalyzed by *A. fulgidus* DsrMKJOP (blue), in the presence of 1mM sulfite (orange), and in the absence of DsrC-trisulfide (purple), and absence of DsrMKJOP (red). Sulfite:DsrC-trisulfide oxidoreductase activity by DsrMKJOP in the absence of menadiol was also followed (grey).
